# GM-CSF orchestrates monocyte and granulocyte responses to *Cryptococcus gattii*

**DOI:** 10.1101/2025.08.01.668083

**Authors:** Alison Ricafrente, Sreemoyee Acharya, Shuyi Chen, Adiza Abass, Aelita Arshakyan, Tyler J. Olson, Lena J. Heung

## Abstract

*Cryptococcus gattii* is an emerging fungal pathogen that is acquired through the respiratory tract and causes invasive infections in both immunocompromised and otherwise healthy people. Many of these apparently immunocompetent patients are subsequently found to have autoantibodies against the pleiotropic cytokine GM-CSF. In this study, we investigated the potential role of GM-CSF (or CSF2) in the host response to *C. gattii* using a murine model of infection. Interestingly, infected *Csf2^-/-^*mice were found to have significantly improved survival and decreased lung fungal burden compared to wild type (WT) mice. We determined that during *C. gattii* infection, GM-CSF promotes the differentiation of monocytes into alveolar and interstitial macrophages. When these macrophages are ablated in CCR2-DTR^+^ mice, there is a corresponding improvement in survival with decreased lung fungal burden, thus phenocopying *Csf2^-/-^* mice. WT bone marrow-derived macrophages challenged with *C. gattii* have poor antifungal activity, suggesting that monocyte-derived macrophages (moMacs) are rendered permissive for fungal proliferation. Therefore, GM-CSF and moMacs mediate immune responses that are harmful to the host. We further found that GM-CSF and moMacs preferentially promote the influx of eosinophils over neutrophils into the infected lung. This eosinophilia is associated with substantial inflammatory lung pathology. Ablation of neutrophils using Mrp8cre^tg^ iDTR^+^ mice significantly increased *C. gattii* burden in the lungs, indicating that GM-CSF and moMacs block the entry of these beneficial, fungal-clearing granulocytes during infection. Altogether, our results show that GM-CSF plays a key role in impeding host anti-fungal responses to *C. gattii* by coordinating monocyte-derived macrophages and granulocyte activity and crosstalk.

**Author Summary:** *Cryptococcus gattii* is an environmental fungus that can cause severe lung and brain infections after inhalation through the respiratory tract. *C. gattii* causes disease in patients with known immune deficits but also in individuals that are apparently healthy. Studies on otherwise healthy people who become infected with *C. gattii* suggest that they may have a previously unrecognized problem involving granulocyte macrophage-colony stimulating factor (GM-CSF), a cytokine, or messenger protein, that is an important part of the immune system. Here, we investigate the role of GM-CSF in the immune response to *C. gattii* using a mouse model of infection. We find that *C. gattii* increases GM-CSF in the lungs, leading to the influx of immune cells, including monocyte-derived macrophages and eosinophils, while inhibiting the entry of neutrophils. The macrophages and eosinophils allow the fungus to proliferate and cause inflammatory damage to the lungs, which is ultimately fatal. The absence of neutrophils also contributes to fungal growth, as these immune cells would otherwise be able to help kill the fungus. Our study provides new insight into how GM-CSF regulates immunity to *C. gattii* and has important implications as to the mechanisms that govern susceptibility to this infection.

## Introduction

*Cryptococcus gattii* is an environmental, encapsulated yeast that is an important cause of invasive lung and brain infections in humans [1]. Known to be endemic in tropical and subtropical parts of the world, *C. gattii* subsequently caused an outbreak of cryptococcosis in 1999 in the temperate regions of British Columbia and the Pacific Northwest, thus broadening its global impact [2–4]. In contrast to its opportunistic relative *Cryptococcus neoformans*, *C. gattii* can also infect apparently healthy individuals. With mortality from *C. gattii* infections estimated between 10-33% [5–7], it is critically important to understand the distinct mechanisms *C. gattii* uses to cause disease in an expanding patient population. Indeed, the World Health Organization designated *C. gattii* one of its first fungal priority pathogens in 2022 given its disease potential and the large knowledge gaps regarding its pathogenicity [8].

Studies on *C. gattii* infections in otherwise healthy people discovered a close correlation with the presence of autoantibodies (AAb) against the cytokine granulocyte macrophage-colony stimulating factor (GM-CSF or CSF2) [9–12]. Anti-GM-CSF AAb are also linked to other fungal and bacterial infections, like aspergillosis and nocardiosis, and the lung disease pulmonary alveolar proteinosis (PAP) [13–15]. GM-CSF is a cytokine that plays a critical role in the development of myeloid cells and their effector functions in a broad range of disease states, from infections to autoimmune disorders [16, 17]. Although *C. gattii* infections are associated with anti-GM-CSF AAb, the role of GM-CSF in the immune response to this pathogen is not well understood.

Here, we used a fatal model of murine *C. gattii* infection to establish that GM-CSF hinders clearance of infection and promotes immunopathology in the lungs. *Csf2*^-/-^ mice lacking GM-CSF have significantly improved survival rates and decreased lung fungal burden compared to WT mice. We found that GM-CSF facilitates the differentiation of CCR2^+^Ly6C^hi^ monocytes into alveolar and interstitial macrophages and that ablating these immune cells in CCR2-DTR^+^ mice phenocopies the improved infectious outcome we observed in *Csf2*^-/-^ mice. Using bone marrow-derived macrophages, we established that monocyte-derived macrophages (moMacs) have a direct role in promoting *C. gattii* proliferation because they are poorly activated when challenged with the fungus. We also determined that GM-CSF and moMacs support the pulmonary infiltration of eosinophils, that cause significant airway inflammation, while also blocking the entry of neutrophils that would otherwise be beneficial for fungal clearance. Together, these results indicate a critical role for GM-CSF in regulating a monocyte-granulocyte axis that determines host outcomes during *C. gattii* infection.

## Results

### GM-CSF mediates poor host outcomes after *C. gattii* infection

To evaluate the role of GM-CSF during *C. gattii* infection, we first established a fatal, respiratory infection model by administering 10^5^ *C. gattii* strain R265 intratracheally (i.t.) to wild type (WT) C57BL/6J mice (S1 Fig). In these WT mice, total lung GM-CSF levels increased after infection, peaking at Day 7 post-infection (p.i.) (Fig 1A). When *Csf2*^-/-^ mice that lack GM-CSF were infected, they had prolonged survival with a median of 43 days as compared to 16 days for WT mice (Fig 1B). *Csf2*^-/-^ mice were able to control fungal proliferation in the lungs, while WT mice had an approximately 1 log increase in lung fungal burden between Days 5 and 10 p.i. (Fig 1C). There were no significant differences in mediastinal lymph node (MLN) or brain fungal burden (Figs 1D and 1E). Grossly, the lungs of WT mice appeared significantly abnormal compared to that of *Csf2*^-/-^ mice, with enlarged, nodular lobes evident by Day 10 p.i. (Fig 1F). On histology, WT mice had noticeably enlarged alveolar spaces at Day 10 p.i. compared to *Csf2*^-/-^ mice (Figs 1G and 1H), and these spaces were replete with proliferating fungal cells (Figs 1I and 1J). We also observed that fungal cells intercalated the collagen fibers within the adventitial sheath between bronchi and arterioles in WT mice (Fig 1K), which was not seen in *Csf2*^-/-^ mice (Fig 1L). Collectively, these data demonstrate that GM-CSF promotes fungal proliferation and invasion and distortion of lung architecture, leading to accelerated mortality rates during *C. gattii* infection.

**Fig 1.**
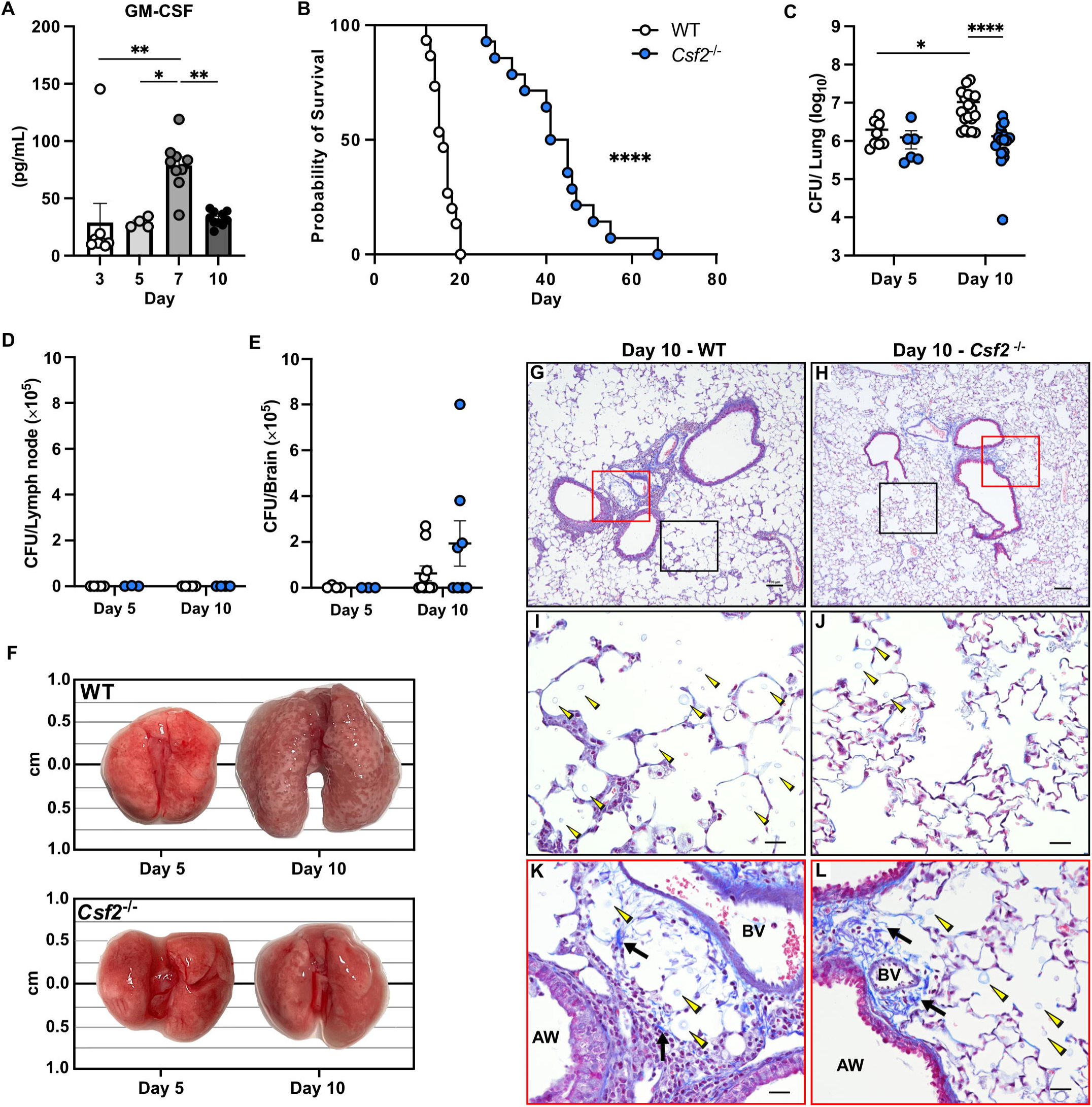
GM-CSF mediates detrimental host responses to *Cryptococcus gattii*. (A) Pulmonary GM-CSF cytokine levels in WT mice infected with *C. gattii*. Days 3, 7, and 10 cytokine data include *n*=8-10 total mice per timepoint from *N*=2 independent experiments, except Day 5 data are from *n*=5 total mice from *N*=1 experiment. (B) Kaplan-Meier survival curve of WT (white circles) and *Csf2^-/-^* (blue circles) mice, *n*=14-15 mice per group from *N*=3 independent experiments. (C-E) Fungal burden was measured in lung (C), mediastinal lymph node (D), and brain (E). For the lung data, Day 5 includes *n*=6-8 total mice per group from *N*=2 independent experiments, and Day 10 includes *n*=16-19 total mice per group from *N*=3 independent experiments. For lymph node and brain data, Day 5 consists of *n*=3-4 total mice per group from *N*=1 experiment, and Day 10 consists of *n*=8-10 total mice per group from *N*=2 independent experiments. (F) Representative images of unaltered whole lungs surgically collected from WT and *Csf2^-/-^* mice at Days 5 and 10 p.i. (G-L) Representative Masson’s trichrome stain of bronchovascular bundles in WT (G) and *Csf2^-/-^* (H) mice at Day 10 p.i., scale bar = 100 μM at 10X magnification. Black box is a magnified inset for the alveolar spaces of WT (I) and *Csf2*^-/-^ (J) mice, and red box is a magnified inset for the adventitial sheath of WT (K) and *Csf2*^-/-^ (L) mice, scale bar = 12 μM at 40X magnification. Airway (AW); Blood vessel (BV); Collagen fibers (black arrows); Fungal cells (yellow arrowheads). Data presented as mean ± SEM and analyzed using one-way ANOVA (A), Mantel-Cox test (B), or two-way ANOVA (C-E). *, *P* <0.05. **, *P* <0.01. ****, *P* <0.0001.

### GM-CSF facilitates monocyte differentiation into effector cells during *C. gattii* infection

Since GM-CSF is known to regulate myeloid cells, we investigated what specific immune cells may be mediating downstream effects. On histology, we observed that cryptococcal cells in WT lungs were surrounded by large foamy macrophages (Fig 2A), while in *Csf2*^-/-^ lungs there was a notable lack of macrophages in similar sites (Fig 2B). *Csf2*^-/-^ mice are known to have a congenital defect in alveolar macrophage development [18, 19], which we confirmed in infected *Csf2*^-/-^ versus WT mice by flow cytometry (Fig 2C). Additionally, interstitial macrophages are significantly reduced in *Csf2*^-/-^ lungs by Day 10 p.i. compared to WT (Fig 2D). During inflammation or infection, alveolar and interstitial macrophages can be replenished by CCR2^+^Ly6C^hi^ monocytes that are recruited to the lungs [20–23]. Despite an increase in CCR2^+^Ly6C^hi^ monocytes in *Csf2*^-/-^ lungs at Day 10 p.i. (Fig 2E), there was no corresponding increase in alveolar or interstitial macrophages (Figs 2C and 2D). CCR2^+^Ly6C^lo^ monocytes, another derivative of CCR2^+^Ly6C^hi^ monocytes [24, 25], are elevated in *Csf2*^-/-^ mice at Day 10 p.i. (Fig 2F), and monocyte-derived dendritic cells (moDCs) are present in comparable numbers in both mouse strains (Fig 2G). These results suggest that the lack of GM-CSF limits the differentiation of recruited monocytes into macrophages, as studies have demonstrated in other infection models [26–28]. Monocyte chemoattractant protein-1 (MCP-1) or CCL2, a chemokine that mobilizes monocytes from the bone marrow, decreases in *Csf2*^-/-^ lungs at Day 10 p.i., suggesting a backlog of monocytes in the lungs (Fig 2H) [29], and decreases in IL-1α and IL-1β in *Csf2*^-/-^ lungs (Figs 2I and 2J) may be attributable to the absence of macrophages [30]. Taken together, these studies suggest that GM-CSF regulation of monocyte differentiation into macrophages may play a key role in the immune response during *C. gattii* infection.

**Fig 2.**
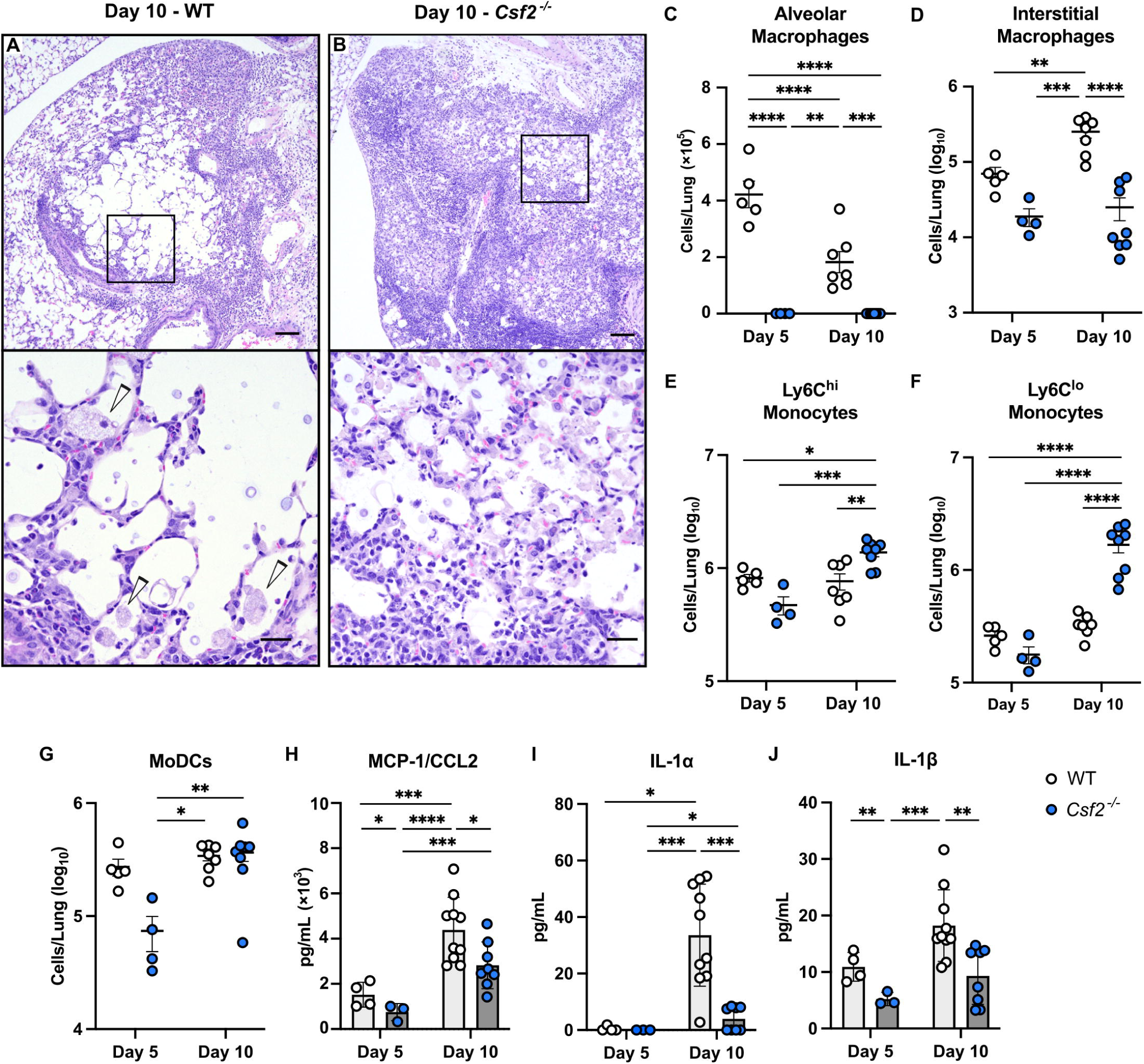
GM-CSF regulates the generation of monocyte-derived macrophages during *C. gattii* infection. (A-B) Representative H&E staining of infected WT (A) and *Csf2^-/-^*(B) lung sections at Day 10 p.i. showing pulmonary infiltrates, scale bar = 200μM at 4X magnification. Black boxes indicate magnified insets in bottom panels, scale bar = 12μM at 60X magnification. Foamy macrophages (white arrowhead). (C-G) Flow cytometry of lung cells from WT (white circles) and *Csf2^-/-^* (blue circles) mice at Days 5 and 10 p.i., including alveolar macrophages (C), interstitial macrophages (D), Ly6C^hi^ monocytes (E), Ly6C^lo^ monocytes (F), and monocyte-derived dendritic cells (moDCs) (G). (H-J) Pulmonary levels of MCP-1/CCL2 (H), IL-1!Z (I), and IL-1β (J). For flow cytometric and cytokine data, Day 5 includes *n*=3-5 total mice per group from *N*=1 experiment; Day 10 includes *n*=7-10 total mice per group from *N*=2 independent experiments. Data presented as mean ± SEM and analyzed using two-way ANOVA (C-J). *, *P* <0.05. **, *P* <0.01. *** *P* <0.001. ****, *P* <0.0001.

### Ablation of monocytes and their derivatives improves host outcomes after *C. gattii* infection

To determine the role of monocytes and their derivative cells during *C. gattii* infection, CCR2-DTR^+^ mice [31] treated with diphtheria toxin (DT) (Fig 3A) were used to ablate CCR2^+^Ly6C^hi^ monocytes, which also resulted in the expected loss of CCR2^+^Ly6C^lo^ monocytes, alveolar and interstitial macrophages, and moDCs compared to WT littermate controls (S2A – S2E Figs). This ablation of monocytes and their derivatives improved host outcomes, as CCR2-DTR^+^ mice had a significantly increased survival rate of 64% as compared to 13% in WT mice (Fig 3B). CCR2-DTR^+^ mice also had a significant reduction in fungi in the lungs at Days 3 and 7 p.i. (Fig 3C) and in the MLN at Day 3 p.i. (Fig 3D) with no significant difference in brain fungal burden (Fig 3E). Increases in GM-CSF and MCP-1/CCL2 in the lungs of CCR2-DTR^+^ mice at Days 3 and 7 p.i. suggest a feedback loop in response to the loss of these cells (S2F and S2G Figs). Similar to *Csf2*^-/-^ mice, the lungs of CCR2-DTR^+^ mice appeared grossly normal at Day 7 p.i. (S2J Fig) with less alveolar enlargement (S2K and S2L Figs) and less fungal invasion of the adventitial sheath (S2M and S2N Figs) compared to WT mice. Thus, the absence of monocytes and their derivatives improves host outcomes after *C. gattii* infection in a similar pattern as when GM-CSF is absent.

**Fig 3.**
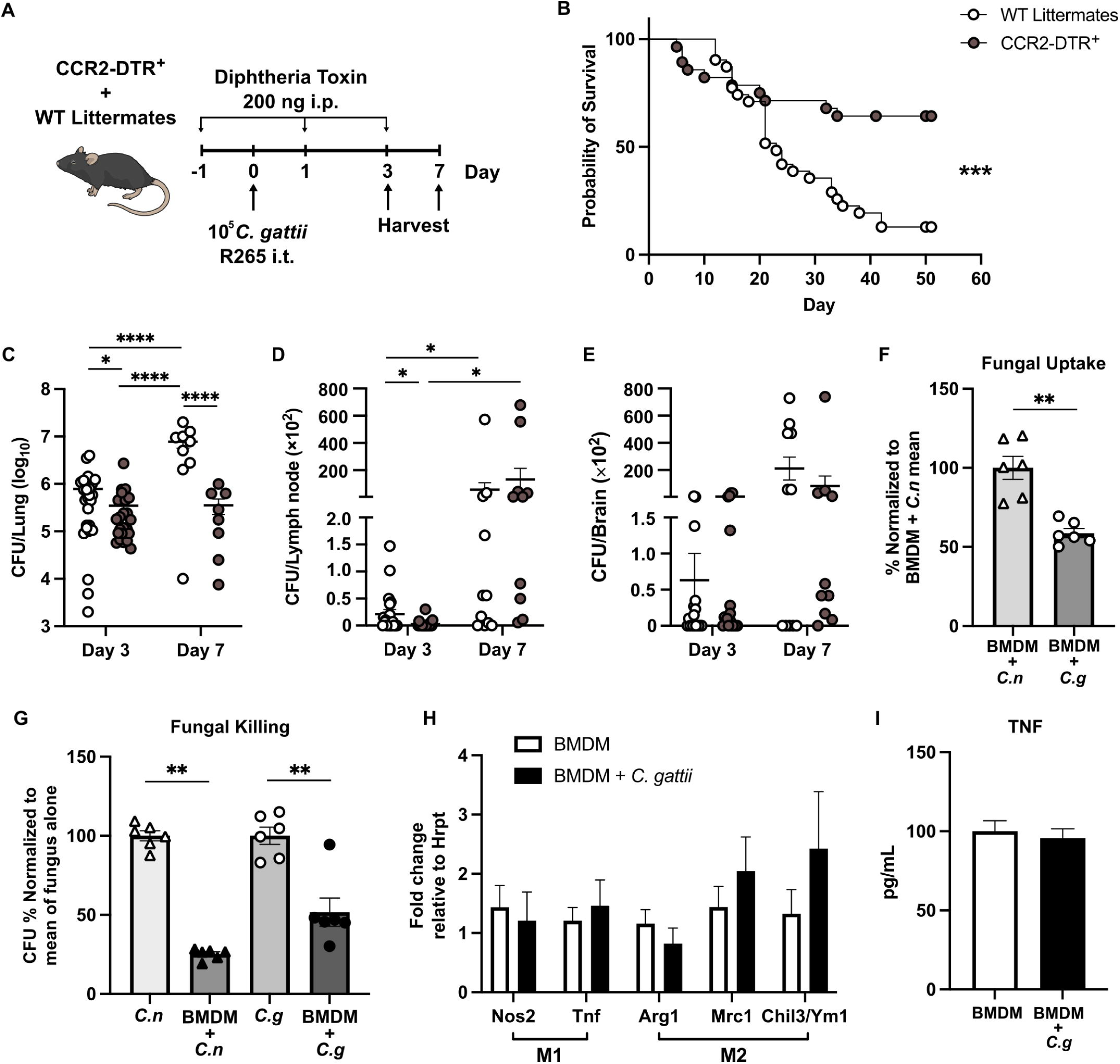
Monocytes and monocyte-derived macrophages are permissive for *C. gattii* infection. (A) Monocyte ablation strategy using CCR2-DTR^+^ and WT littermate mice, treated with 200 ng of diphtheria toxin by i.p. injection a day prior to infection with *C. gattii* and at Days 1 and 3 p.i. Organs were harvested at Day 3 or 7 p.i. for fungal burden measurements. (B) Kaplan-Meier survival curve of WT littermate (white circles) and CCR2-DTR^+^ (gray circles) mice, *n*=28-31 total mice per group from *N*=6 independent experiments. (C-E) Fungal burden was measured in lung (C), MLN (D), and brain (E). For lungs, Day 3 includes *n*=24-26 total mice per group from *N*=6 independent experiments; Day 7 includes *n*=8-9 total mice per group from *N*=2 independent experiments. For MLN and brains, Day 3 consists of *n=*18-21 total mice per group from *N*=5 independent experiments; Day 7 consists of *n=*10-11 total mice per group from *N*=2 independent experiments. (F-I) BMDM were challenged with 2×10^4^ *C. neoformans* or *C. gattii* at an MOI of 1:40 and analyzed for fungal uptake (F), fungal killing (G), expression of M1 and M2 polarization markers by qRT-PCR (H), and TNF cytokine secretion (I). Results are from *N=*2-3 independent experiments with *n*=6-9 total replicates per group. Data presented as mean ± SEM and analyzed using Mantel-Cox test (B), two-way ANOVA (C-E), and Mann-Whitney test (F-I). *, *P* <0.05. **, *P* <0.01. ***, *P* <0.001. ****, *P* <0.0001.

### Monocyte-derived dendritic cells do not mediate immune responses to *C. gattii*

To evaluate if moDCs play any role in determining outcomes after *C. gattii* infection, we generated CCR2-Cre^+^ MHCII^fl/fl^ mice. In this mouse model, both moDCs and conventional DCs (cDCs) lack MHCII for antigen presentation compared to MHCII^fl/fl^ littermate controls (S3A Fig), as cDCs are reported to express CCR2 [32, 33]. Despite the loss of MHCII expression by these DC subsets, along with a reduction in total numbers of moDCs (S3B Fig), we found no differences in survival or organ fungal burden in CCR2-Cre^+^ MHCII^fl/fl^ mice versus controls after *C. gattii* infection (S3C – S3F Figs). These data indicate that moDCs and cDCs do not play a critical role in the immune response to *C. gattii*. Thus, observed reductions in cDCs in *Csf2*^-/-^ mice [34] (S4G and S4H Figs) and in moDCs and cDCs in CCR2-DTR^+^ mice (S2E, S3I, and S3J Figs) are unlikely to have contributed significantly to the phenotypes observed. Rather, monocyte-derived alveolar and interstitial macrophages appear to facilitate the progression of disease in our model of *C. gattii* infection.

### Monocyte-derived macrophages are permissive for *C. gattii* proliferation

To study the direct role of monocyte-derived macrophages (moMacs) in the immune response to *C. gattii*, bone marrow-derived macrophages (BMDM) from WT mice were challenged with *C. gattii* R265-GFP or *C. neoformans* H99-GFP [35] as a comparator. We previously established that moMacs are subverted by *C. neoformans* to enable fungal proliferation in the lungs [36]. When BMDM were challenged with *C. gattii*, fungal uptake of *C. gattii* by BMDM was significantly reduced by an average of 41% relative to the uptake seen with *C. neoformans* (Fig 3F). Although there was some killing of both *C. gattii* and *C. neoformans* in the presence of BMDM, the killing of *C. gattii* by BMDM was significantly impaired, with only a 48% reduction in *C. gattii* colony forming units (CFUs) relative to fungus alone as compared to a 75% reduction in *C. neoformans* CFUs relative to fungus alone (Fig 3G). Interestingly, *C. gattii* does not induce significant polarization of BMDM since there were no changes in the expression of typical markers for M1 (*Nos2* and *Tnf*) or M2 (*Arg1*, *Mrc1* and *Ym1*/*Chil3*) macrophage polarization (Fig 3H). There was also no change in TNF secretion by *C. gattii*-infected versus non-infected BMDM (Fig 3I). These results demonstrate that moMacs are unable to effectively respond directly to *C. gattii* infection, providing an environment that allows for fungal proliferation.

### GM-CSF and monocyte-derived macrophages induce pulmonary eosinophilia

When comparing our findings in *Csf2*^-/-^ and CCR2-DTR^+^ mice, we observed similar patterns of granulocyte infiltration into the lungs, suggesting GM-CSF and moMacs may further regulate host outcomes through crosstalk with these immune cells. On lung histology there was a predominance of eosinophils in perivascular infiltrates in WT controls (Figs 4A and 5A), while in *Csf2*^-/-^ and CCR2-DTR^+^ mice there was a clear shift towards neutrophil infiltration (Figs 4B and 5B). Flow cytometry validated that total lung eosinophils were significantly higher in WT controls than in *Csf2*^-/-^ and CCR2-DTR^+^ mice at all timepoints (Figs 4C and 5C). The relative numbers of total lung neutrophils were more variable when comparing WT controls and *Csf2*^-/-^ and CCR2-DTR^+^ mice over time (Figs 4D and 5D). However, as infection progressed, WT controls consistently demonstrated a higher eosinophil-to-neutrophil ratio in the lungs compared to either *Csf2*^-/-^ or CCR2-DTR^+^ mice (Figs 4E and 5E). We evaluated whole lung cytokines and chemokines that might facilitate these differences in granulocyte recruitment. A key finding was that the eosinophil-associated cytokine IL-5 is significantly increased in the lungs of WT controls compared to *Csf2*^-/-^ and CCR2-DTR^+^ mice (Figs 4F and 5F). Conversely, the neutrophil-associated cytokine granulocyte-colony stimulating factor (G-CSF) is suppressed in WT lungs versus *Csf2^-/-^* and CCR2-DTR^+^ mice (Figs 4G and 5G). There is an increase in the cytokine subunit IL-12p40 in WT controls (Figs 4H and 5H), without any change in its Th1-associated heterodimer IL-12p70 (Figs 4I and 5I) or other Th1 and inflammatory cytokines (S2H-S2I, S4A and S5A Figs) as compared to *Csf2^-/-^* and CCR2-DTR^+^ mice. WT controls also demonstrated a skew towards higher amounts of additional Th2-associated cytokines and chemokines, most prominent in comparison to CCR2-DTR^+^ mice (S4B-S4C and S5B-S5C Figs), and an increase in the Th17 cytokine IL-17 (Figs 4J and 5J). These pro-eosinophilic conditions support clear evidence of airway inflammation in WT lungs, including goblet cell hyperplasia and metaplasia in the small and terminal airways and thickening of the subepithelial matrix with increased collagen deposition that is not present in *Csf2^-/-^* and CCR2-DTR^+^ mice (S6 Fig). These findings suggest that GM-CSF and moMacs modulate cytokines and chemokines to preferentially induce pulmonary eosinophilia and inflammatory lung damage after *C. gattii* infection.

**Fig 4.**
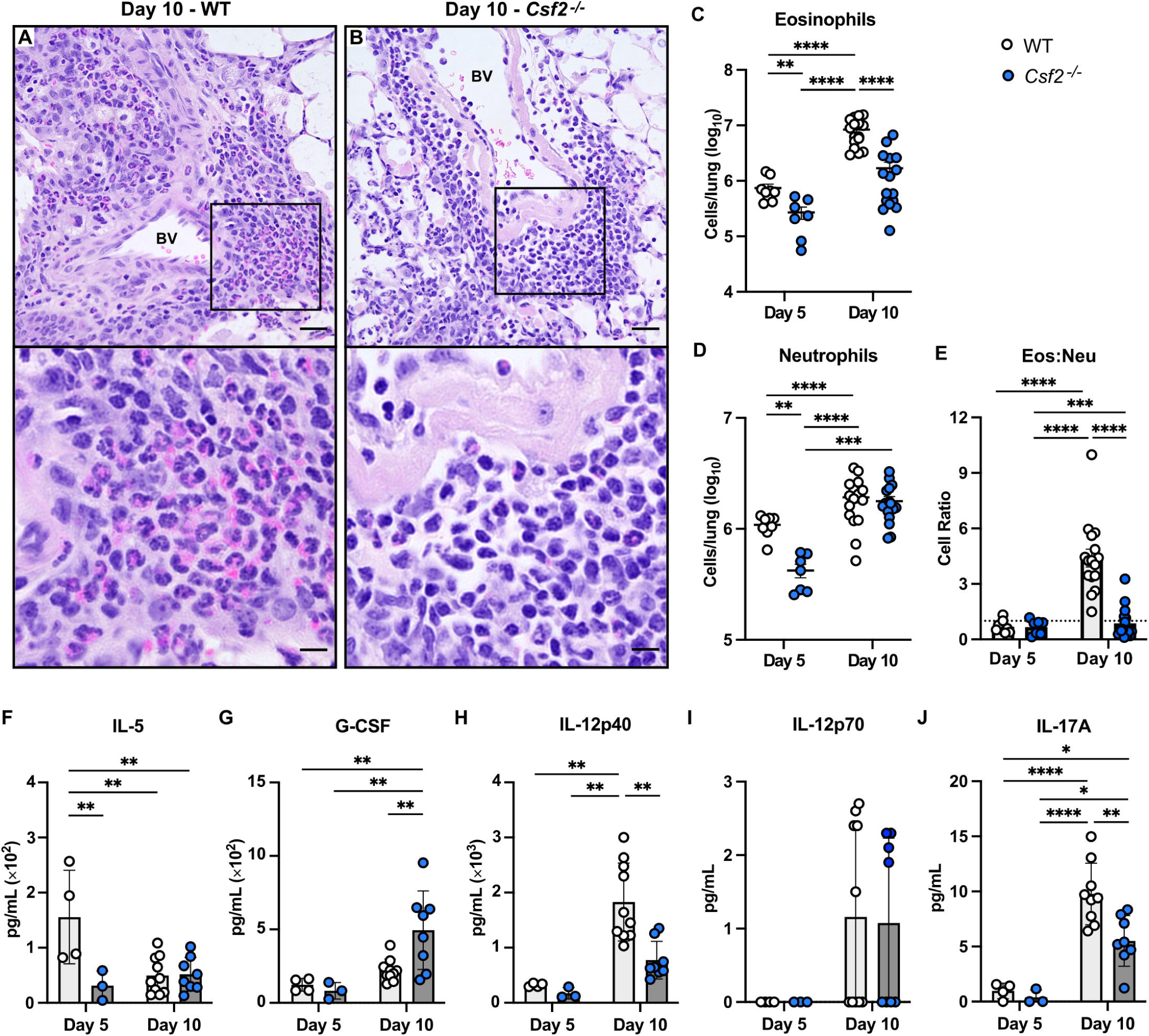
GM-CSF induces the pulmonary influx of eosinophils over neutrophils during infection. (A-B) Representative H&E staining of lung sections from WT (A) and *Csf2^-/-^*(B) mice at Day 10 p.i. showing perivascular cell infiltrates, scale bar = 25μM at 20X magnification. Black boxes represent magnified insets in bottom panels, scale bar = 12μM at 60X magnification. BV (Blood vessel). (C-E) Flow cytometry of lung cells from WT (white circles) and *Csf2^-/-^*(blue circles) mice, including eosinophils (C) neutrophils (D) and the calculated eosinophil-to-neutrophil (Eos:Neu) cellular ratio (E); a ratio of 1:1 is indicated by the dotted line. Day 5 is *n*=7-9 total mice per group from *N*=2 independent experiments; Day 10 is *n*=16 mice per group from *N*=4 independent experiments. (F-J) Pulmonary levels of cytokines IL-5 (F), G-CSF (G), IL-12p40 (H), IL-12p70 (I), and IL-17 (J). Day 5 is *n*=3-10 total mice per group from *N*=1-2 independent experiments; Day 10 is *n=*8-10 total mice per group from *N=*2 independent experiments. Data presented as mean ± SEM and analyzed using two-way ANOVA (C-J). *, *P* < 0.05. **, *P* < 0.01. ***, *P <* 0.001. ****, *P* <0.0001.

**Fig 5.**
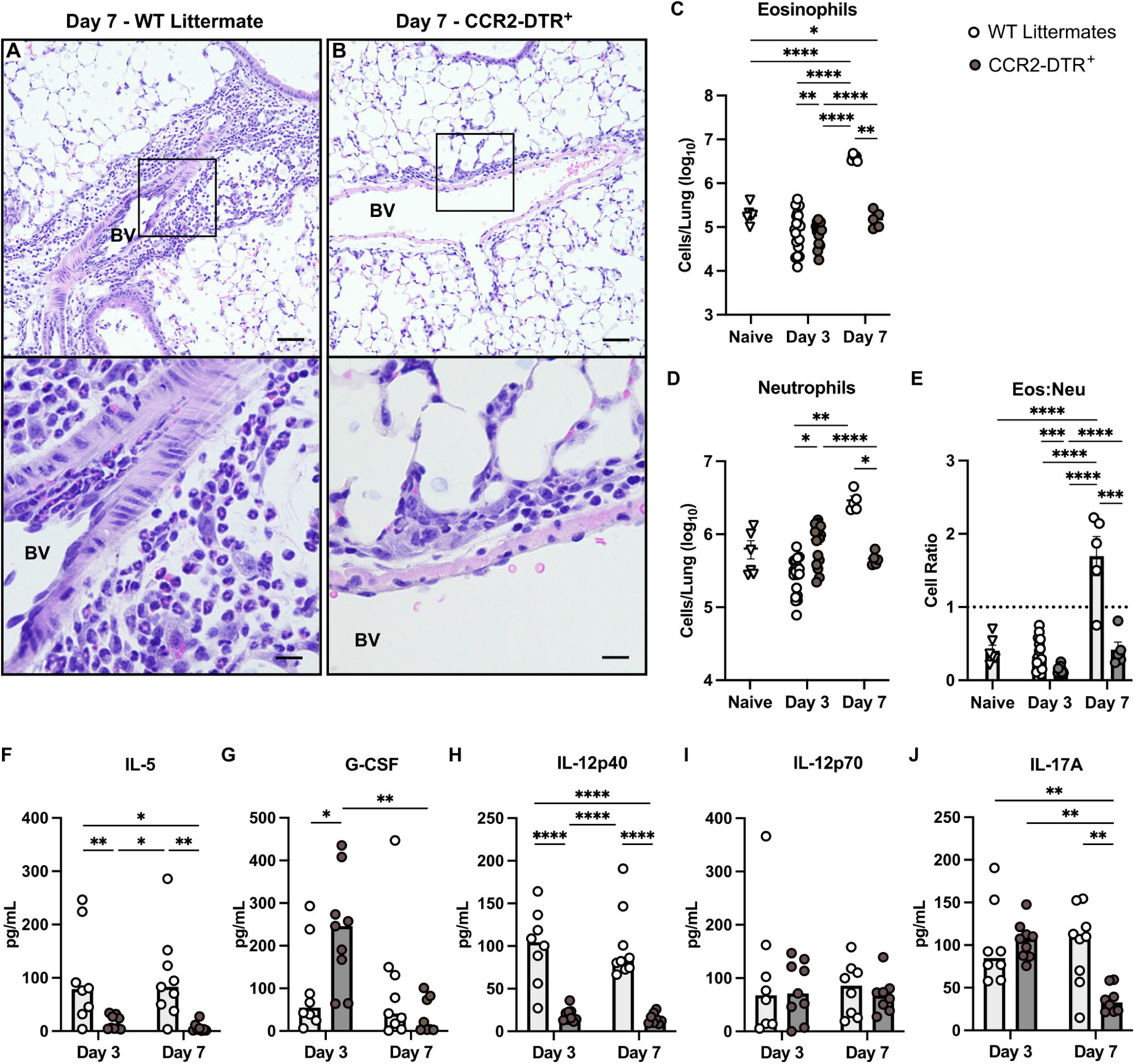
Monocytes and monocyte-derived macrophages reinforce a high eosinophil-to-neutrophil ratio in the infected lung. (A-B) Representative H&E staining of lung sections from WT littermates (A) and CCR2-DTR^+^ (B) mice at Day 7 p.i. showing perivascular cell infiltrates, scale bar = 25μM at 20X magnification. Black boxes represent magnified insets in bottom panels, scale bar = 12μM at 60X magnification. BV (Blood vessel). (C-E) Flow cytometry of lung cells from naive WT (white triangles) and infected WT littermates (white circles) and infected *Csf2^- /-^* mice (gray circles), including eosinophils (C), neutrophils (D), and the calculated Eos:Neu cellular ratio (E); a ratio of 1:1 is indicated by the dotted line. Results include *n*=6 naive WT littermates from *N=*2 independent experiments and *n=*5-19 total infected mice per group per timepoint from *N*=1-3 independent experiments. (F-J) Pulmonary levels of cytokines IL-5 (F), G-CSF (G), IL-12p40 (H), IL-12p70 (I), and IL-17 (J). Results are from *n*=8-9 total mice per group per timepoint from *N*=2 experiments. Data presented as mean ± SEM and analyzed using two-way ANOVA (C-J). *, *P* < 0.05. **, *P* < 0.01. ***, *P <* 0.001. ****, *P <* 0.0001.

### GM-CSF and monocyte-derived macrophages suppress neutrophil influx to promote *C. gattii* proliferation

To determine if the reduction in pulmonary neutrophils observed in WT control mice affects progression of *C. gattii* infection, we utilized Mrp8cre^tg^ iDTR^+^ mice to deplete these cells by i.p. injection of diphtheria toxin (Fig 6A) [37]. The Mrp8cre^tg^ iDTR^+^ mice were infected alongside iDTR^+^ littermate controls, and then whole lungs were collected for analysis. We observed an approximately 90% (1 log) decrease in lung neutrophils in Mrp8cre^tg^ iDTR^+^ mice at Day 7 p.i. compared to controls (Fig 6B). This loss of neutrophils in Mrp8cre^tg^ iDTR^+^ mice resulted in a significant increase in lung fungal burden versus control mice (Fig 6C), suggesting neutrophils are beneficial for limiting *C. gattii* growth in the lungs. Thus, the absence of antifungal neutrophils in combination with the presence of permissive moMacs and pulmonary eosinophilia, all driven by GM-CSF, provides an optimal environment for fungal proliferation and lung damage resulting in host mortality.

**Fig 6.**
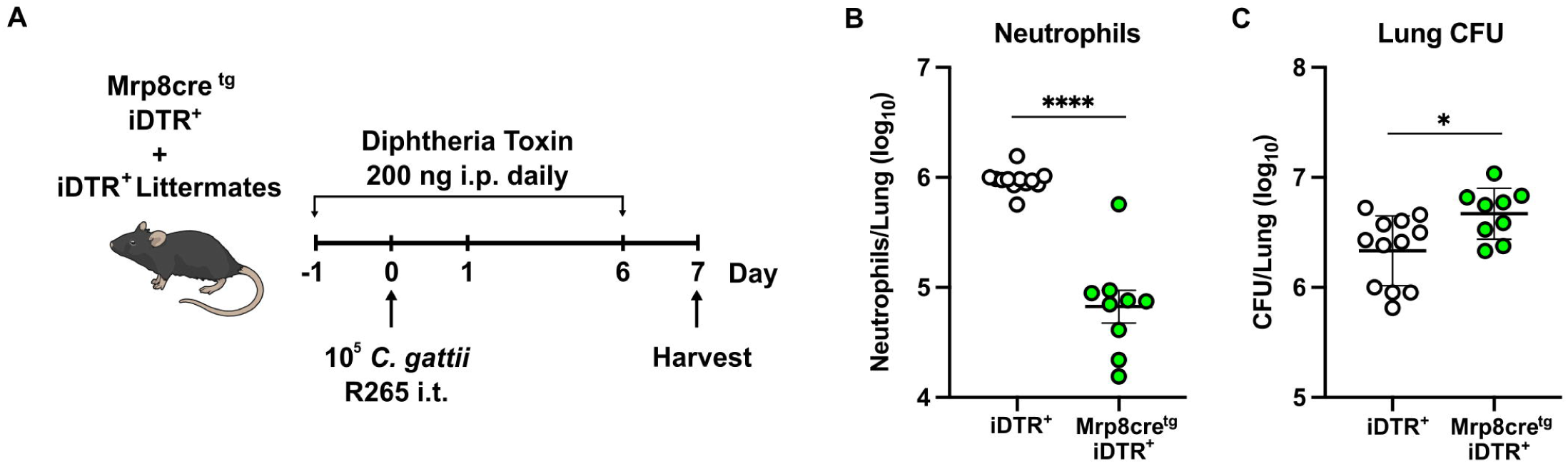
Neutrophils are beneficial to the host after *C. gattii* infection. (A) Neutrophil ablation strategy using Mrp8cre^tg^ iDTR^+^ mice and iDTR^+^ littermate control mice. All mice were treated with 200 ng of diphtheria toxin by i.p. injection daily, starting one day prior to i.t. infection with *C. gattii*. At Day 7 p.i., whole lungs were collected to measure total neutrophils by flow cytometry (B) and fungal burden by CFU (C). Results are from *n*=9-12 total mice per group from *N*=3 independent experiments. Data presented as mean ± SEM and analyzed using Mann-Whitney test. *, *P* < 0.05. ****, *P* <0.0001.

## Discussion

Although *C. gattii* was recognized as an emerging fungal pathogen over 25 years ago [38–40], our understanding of its interactions with the host immune system that lead to infection remains disproportionally limited. In this study, we establish that *C. gattii* induces GM-CSF signaling in the lungs, resulting in poor infectious outcomes. Our findings contrast with a recent study in which administration of recombinant murine GM-CSF led to decreased amounts of *C. gattii* in the lungs [41]. However, that study used a non-fatal infection model with BALB/c mice that are inherently more resistant to *C. gattii* than the C57BL/6J mice used in our work [41, 42]. We instead found that physiologically elevated levels of GM-CSF in the lungs of C57BL/6J mice are associated with significantly increased *C. gattii* burden, immunopathology, and accelerated mortality rates. Thus, GM-CSF may play different roles in acute versus chronic infection, as has also been observed with *C. neoformans* [27, 43–46].

Based on our studies, GM-CSF facilitates the differentiation of recruited CCR2^+^Ly6C^hi^ monocytes into alveolar and interstitial macrophages after *C. gattii* infection. *Cryptococcus* species are facultative intracellular pathogens [1], so these moMacs may be providing a reservoir for fungal cells. Indeed, we observed that BMDM were less able to control *C. gattii* as compared to *C. neoformans*; in previous work, we established that moMacs are already very permissive for *C. neoformans* proliferation [36]. Interestingly, *C. gattii* has reduced uptake by BMDM compared to *C. neoformans*. However, clinical isolates of *C. gattii* were shown to have increased intracellular proliferation in macrophages despite lower phagocytosis rates when measured alongside *C. neoformans* [47]. BMDM also do not seem to polarize significantly and exhibit no change in TNF secretion in response to *C. gattii*. Previous studies have highlighted the immune silencing capacity of *C. gattii* that includes inhibition of genes associated with macrophage autophagy and polarization [48]. Thus, *C. gattii* can efficiently take advantage of moMacs generated under the direction of GM-CSF.

Granulocytes also appear to have a significant role downstream of GM-CSF and MoMacs during infection. The worse infectious outcomes in WT mice, as compared to both *Csf2*^-/-^ and CCR2-DTR^+^ mice, are associated with an increase in eosinophils and a decrease in neutrophils in the lungs. Eosinophils have well-established roles in a wide range of inflammatory pulmonary disorders like asthma and COPD [49–51]. Accordingly, the pulmonary eosinophilia we see in WT mice is associated with significant airway remodeling including goblet cell hyperplasia and metaplasia and collagen deposition in a thickened subepithelial matrix, similar to that described in asthma [52, 53]. GM-CSF is known to promote the accumulation and survival of eosinophils in allergic airway inflammation [54], and macrophages can play a role in recruiting eosinophils to peripheral tissues [55]. We also see increases in other cytokines in WT mice that can promote eosinophils, including IL-5, IL-12p40, and IL-17. IL-5 has established roles in supporting the development, accumulation, and activity of eosinophils [54, 56]. IL-12p40 can be secreted by macrophages and monocytes [57], and as a free monomer or homodimer, IL-12p40/IL-12p80 can act as an antagonist of IL-12 signaling [58–61]. IL-12 signaling could otherwise induce apoptosis by eosinophils and promote antimicrobial neutrophil activity [62, 63]. IL-17 (or IL-17A) is generally known to drive neutrophil responses [64], but it can also induce pulmonary eosinophilia during allergic aspergillosis [65] and be further secreted by eosinophils themselves [66]. The absence of neutrophils in the infected WT lung is likely due, in part, to the observed decreased levels of G-CSF, a cytokine that typically promotes neutrophil migration and activity in peripheral tissues, in addition to its role in granulopoiesis [67]. This lack of neutrophil influx is significant, as we demonstrate that neutrophil-depleted mice are unable to control *C. gattii* proliferation in the lungs. Previous in vitro studies have also found that mouse and human neutrophils have fungicidal activity against *C. gattii* [41, 68]. In sum, GM-CSF may have direct and indirect roles, via moMacs and other immune cells, in the control of granulocyte influx and subsequent pathology that develops during *C. gattii* infection (Fig 7).

**Fig 7.**
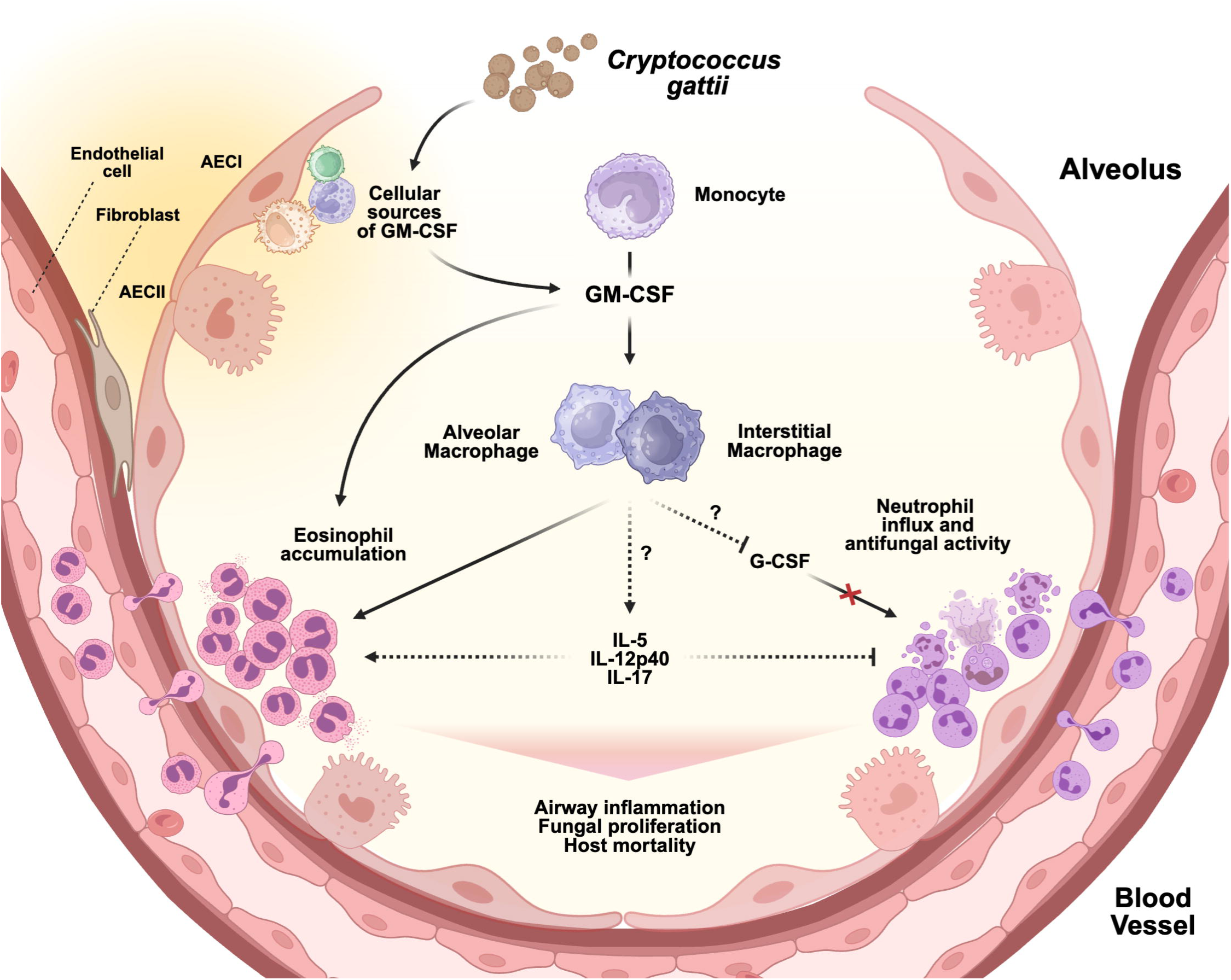
Schematic model of GM-CSF and monocyte signaling during *Cryptococcus gattii* infection. After *C. gattii* enters the lungs, the fungus may induce various cells, such as Type I (AECI), Type II alveolar epithelial cells (AECII), endothelial cells, or fibroblasts to release GM-CSF. This GM-CSF induces recruited CCR2^+^Ly6C^hi^ monocytes to differentiate into alveolar and interstitial macrophages. However, these macrophages remain in an unpolarized M0 state, which enables *C. gattii* to proliferate and invade host tissues unchecked. GM-CSF and monocyte-derived macrophages also facilitate the pulmonary influx of eosinophils while inhibiting the entry of neutrophils through the regulation of cytokines including IL-5, G-CSF, IL-12p40, and IL-17. The accumulation of eosinophils leads to harmful airway inflammation. The absence of antifungal neutrophils allows further fungal proliferation. Altogether, the secretion of GM-CSF and the generation of monocyte-derived macrophages fuel an imbalance of eosinophils and neutrophils that ultimately results in host mortality. Created in BioRender. Ricafrente, A. (2025) https://BioRender.com/lpowwec

We recognize that our study has only addressed some facets of how GM-CSF may regulate *C. gattii* infection, and it remains to be investigated if these mechanisms are recapitulated in human models. Although GM-CSF is linked to beneficial host responses in other fungal diseases [27, 43–45, 69–72], there is some precedent for GM-CSF increasing susceptibility to infections. In particular, pulmonary GM-CSF has been linked to worse clinical outcomes for severe COVID-19 due to its role in dysregulation of myeloid cells [73, 74]. This observation has led to trials on the use of anti-GM-CSF monoclonal antibodies as potential treatments for COVID [75, 76]. While our work did not directly study the role of anti-GM-CSF AAb, our findings challenge the prevailing idea that these autoantibodies increase susceptibility to *C. gattii* [12, 77]. Thus, it remains a possibility that anti-GM-CSF AAb do not play a pathogenic role in cryptococcosis, which could explain why their serum levels or degree of neutralization activity do not seem to have a direct correlation to the development of disease [12, 78]. Rather, the induction of GM-CSF by *C. gattii* may set up conditions for autoimmunization, resulting in the development of anti-cytokine autoantibodies which can be a physiologic process in healthy people to help maintain homeostasis [79, 80]. We also note that although the phenotypes of *Csf2*^-/-^ and CCR2-DTR^+^ mice are very similar, some features do not fully overlap and, thus, indicate the presence of mechanisms that may be regulated by GM-CSF or monocytic cells independently. For example, CCR2-DTR^+^ mice exhibit sustained survival during the observation period while *Csf2*^-/-^ mice only have delayed mortality, suggesting monocytes and their derivatives have a more encompassing impact on infectious outcomes than GM-CSF alone. Additionally, CCR2-DTR^+^ mice had more profound decreases in Th2-associated cytokine responses in the lungs compared to *Csf2*^-/-^ mice. Future work using targeted, cell-specific approaches will help to further define the attributable downstream effects of GM-CSF versus monocytes and to identify additional cells and signaling molecules that may facilitate their roles in host immunity.

While we still have much to learn about how *C. gattii* causes disease, our study provides additional insight into the key cell types and signaling pathways that are critical to host susceptibility. We have established a new role for GM-CSF signaling in the pathogenesis of *C. gattii* through its regulation of moMacs and crosstalk with granulocytes during cryptococcosis. This work lays a foundation for further investigation of GM-CSF, monocytes, and moMacs as potential immunomodulatory targets to reduce mortality and other complications from *C. gattii* infection.

## Materials and Methods

### Preparation of Cryptococcus

*C. gattii* strain VGIIa R265 (MYA-4093) was obtained from ATCC. *C. neoformans* strain H99 #4413 was kindly provided by Joseph Heitman (Duke University). Fluorescent strains R265-GFP and H99-GFP were a generous gift from Robin May (University of Birmingham)[35]. All *Cryptococcus* strains were grown on Sabouraud dextrose agar (SAB) plates from frozen glycerol stocks and then cultured overnight at 37°C in YPD broth (1% yeast extract, 2% peptone, 2% dextrose). Fungal cells were washed three times with sterile phosphate-buffered saline (PBS) and resuspended in PBS for further use.

### Mice

Mice were used at 6-8 weeks of age, unless otherwise noted. C57BL/6J (strain #000664), B6.129X1-*H2-Ab1^b-tm1Koni^*/J (strain #013181, referred to as MHCII^fl/fl^), and *Csf2^-/-^* (strain #026812) mice were purchased from Jackson Laboratory. MRP8-Cre-ires/GFP^+^ ROSA26iDTR^+^ mice (referred to as Mrp8cre^Tg^ iDTR^+^ mice) were generated using Jackson strains #021614 and #007900 and were generously provided by Keith Chan (Houston Methodist). The CCR2-DTR^+^ and CCR2-Cre^+^ mice were generated as previously described [31, 36]. All mouse strains were bred and housed in the Cedars-Sinai Medical Center vivarium under specific pathogen-free conditions. Experiments using CCR2-DTR^+^, CCR2-Cre^+^ MHCII^fl/fl^, and Mrp8cre^Tg^ iDTR^+^ mice included littermate control mice that were weaned from the same litters and co-housed. The genotypes of CCR2-DTR^+^ mice were validated by the presence of CD45^+^CD11b^+^CFP^+^ cells in tail vein blood samples using flow cytometry as previously described [31]. The genotypes of Mrp8cre^Tg^ mice were validated by the presence of CD11b^+^Ly6G^+^GFP^+^ cells in tail vein blood samples using flow cytometry. The genotypes of CCR2-Cre^+^, MHCII^fl/fl^, and ROSA26iDTR^+^ (referred to as iDTR^+^) mice were validated by PCR of ear DNA as previously described [36] or as per Jackson Laboratory protocol.

### Ethics statement

All animal procedures were performed with approval by the Institutional Animal Care and Use Committee at Cedars-Sinai Medical Center under protocol 8429 and were compliant with all applicable provisions established by the Animal Welfare Act and the Public Health Services Policy on the Humane Care and Use of Laboratory Animals.

### Murine infection with *Cryptococcus gattii*

Overnight YPD broth cultures of *C. gattii* R265 were washed three times with sterile PBS and resuspended at a concentration of 10^5^ cells per 50 μL volume. Mice were anesthetized with inhaled isoflurane and 50 μL of the fungal cell suspension was administered intratracheally (i.t.) using a blunt ended 20-gauge dispensing tip, as previously described [81].

### Immune cell ablation

To ablate monocytes and neutrophils, respectively, CCR2-DTR^+^ and Mrp8cre^Tg^ iDTR^+^ mice with their control littermates were injected intraperitoneally (i.p.) with 200 ng (10 ng/g body weight) of diphtheria toxin (List Biological Laboratories) with the frequency indicated in Fig 3A and Fig 6A.

### Fungal burden in organs

After infection, whole organs including lungs, mediastinal lymph nodes (MLN), and brains were collected from euthanized mice into sterile PBS. Lungs and brain were homogenized in gentleMACS™ C tubes using a gentleMACS™ Octo Dissociator (Miltenyi Biotec). Specifically, lung samples were homogenized in 5 mL PBS + 0.5% Bovine Serum Albumin (BSA) (Fisher Scientific) + 2.31 mg/mL Collagenase Type 4 (Worthington Biochemical Corporation) + 100 mg/mL DNAse I grade II (Roche) using the “Mouse Spleen 1” program, incubated at 37°C for 45 min, and further homogenized using the “Mouse Lung 2” program. Brain samples were homogenized in 1 mL PBS using the “Mouse Lung 2” program. MLN were mechanically dissociated using ground glass slides in 1 mL PBS. Tissue homogenates were serially diluted in PBS and cultured on SAB plates at 37°C. After 3 days, fungal colonies were counted and total colony forming units (CFU) per organ were calculated.

### Cytokine and chemokine measurement

To analyze cytokine and chemokine levels, whole mouse lungs were collected into 2 mL PBS containing 1X Halt™ Protease Inhibitor Cocktail (Thermo Fisher Scientific) followed by mechanical tissue dissociation using gentleMACS™ C tubes and the “RNA 1” program on a gentleMACS™ Octo Dissociator (Miltenyi Biotec). Tissue homogenates were centrifuged at 1500 rpm to remove cell debris, and the supernatant was collected. Supernatants were processed using the Bio-Plex Pro Mouse Cytokine 23-Plex Assay (Bio-Rad) and analyzed on a Bio-Plex 200 System (Bio-Rad).

### Flow cytometry

For flow cytometry analysis of whole lungs, single cell suspensions were generated using mechanical tissue dissociation and digestion as described in the “Fungal burden in organs” section of the Materials and Methods. Isolated cells were treated with 1X RBC lysis buffer for 5 min and then washed with 0.5% BSA in PBS before being passed through a 100 μM cell strainer. Total cells were counted using a hemocytometer and then stained with fluorescent antibodies (S1 Table). Flow cytometry data was collected on a LSRFortessa (BD Biosciences) and analyzed with FlowJo software v10.10. Gating strategies are shown in S7-S9 Figs.

### Histopathology

The lungs of euthanized mice were instilled with 4% paraformaldehyde (PFA) in PBS *in situ* via a catheter inserted through an incision in the trachea. The lungs were then harvested and fixed by immersion in 4% PFA overnight and stored in 70% ethanol until processing. Lungs were embedded in paraffin using an ASP300S tissue processor (Leica), and 5 μm sections were generated using a Microm HM 325 microtome (Thermo Fisher Scientific). Lung sections were stained with StatLab™ Masson’s Trichrome stain kit (Fisher Scientific) or Harris Hematoxylin Solution, Modified (Millipore Sigma) and Eosin Y solution, Alcoholic (Millipore Sigma) (H&E). Images were captured using a Nikon Ti2 microscope. Figure images shown of WT and *Csf2*^-/-^ stained lung sections at Day 10 p.i. represent *n*=4-5 mice per group from *N*=1 experiment. Figure images shown of WT littermate and CCR2-DTR^+^ stained lung sections at Day 7 p.i. represent *n*=5 mice per group from *N*=1 experiment.

### Bone-marrow derived macrophage studies

Bone-marrow derived macrophages (BMDM) were generated from C57BL/6J mouse bone marrow using L929 cell-conditioned medium, as previously described [82]. Freshly cultured R265-GFP and H99-GFP cells were opsonized for 1 hr at room temperature with murine anti-glucuronoxylomannan monoclonal antibody 18B7, kindly provided by Arturo Casadevall (Johns Hopkins), at a concentration 10 ug/mL in DMEM media. Opsonized fungal cells were used to challenge BMDM at a multiplicity of infection (MOI) of 1:40 for 24 hours at 37°C 5% CO_2_. Killing of fungal cells by BMDM was measured by plating CFU from BMDM lysed in water. Uptake of the fluorescent fungal cells by BMDM was measured by flow cytometry analysis on an LSRFortessa (BD Biosciences). Mouse TNF cytokine secretion was measured by ELISA (Invitrogen) on cell culture supernatant using a Varioskan Lux Multimode Microplate Reader (Thermo Fisher Scientific). For quantitative RT-PCR, total RNA was extracted from BMDM using TRIzol LS (Thermo Fisher Scientific), and cDNA was generated using a High Capacity RNA to cDNA Kit (Applied Biosystems). The cDNA was inspected and normalized using a Nanodrop Spectrophotometer (Thermo Fisher Scientific), and qRT-PCR was performed on a ViiA 7 Real-Time PCR System (Applied Biosystems) using TaqMan Fast Advanced Master Mix and TaqMan Gene Expression Assays (Thermo Fisher Scientific), including Arg1 (Mm00475988_m1), Mrc1 (Mm01329362_m1), Retnla/Fizz1 (Mm00445109_m1), Hprt (Mm03024075_m1), Nos2 (Mm00440502_m1), Tnf (Mm00443258_m1), and Chil3/Ym1 (Mm00657889_mH). Relative expression of transcripts was normalized using Hprt as a housekeeping gene.

### Statistical analysis

All results were expressed as mean ± SEM. A Mann-Whitney U test was used for statistical analysis of two group comparisons, one-way ANOVA was used for 3 or more infected mouse groups, and two-way ANOVA was used for 3 or more naive and infected mouse groups, unless otherwise noted. Survival data was analyzed by Mantel-Cox test. All statistical analyses were performed with GraphPad Prism software, v10.2.0. A *P* value < 0.05 was considered significant and indicated with an asterisk.

## Acknowledgments

We thank Fayyaz Sutterwala, Suzanne Cassel, and Apurwa Trivedi for helpful comments and all our colleagues in the Women’s Guild Lung Institute and Research Division of Immunology, Keith Chan, and Fotis Nikolos for generously sharing materials, equipment, and their expertise. We acknowledge assistance from the Cedars-Sinai Flow Cytometry Shared Resource and Biobank and Research Pathology Resource.

## Supporting Information

**S1 Fig.** Kaplan-Meier survival curve of wild type mice infected with different inocula of *Cryptococcus gattii* R265 intratracheally. Experiments were terminated on Day 61 post-infection. Data are from *n*=6-7 total mice per group from *N*=2 independent experiments. *, *P* < 0.05. ***, *P* < 0.001.

**S2 Fig.** Effects of monocyte ablation on derivative cells, pulmonary cytokines, and lung pathology. (A-E) Flow cytometry of lung cells from naive and infected WT littermates and infected CCR2-DTR^+^ mice, including Ly6C^hi^ monocytes (A), Ly6C^lo^ monocytes (B), alveolar macrophages (C), interstitial macrophages (D), and moDCs (E). Data include *n*=6 total naive mice from *N=*2 independent experiments; Day 3 is *n*=15-19 total mice per infected group from *N*=3 independent experiments; and Day 7 is *n*=5 total mice per infected group from *N*=1 experiments. (F-G) Pulmonary levels of GM-CSF (F), MCP-1/CCL2 (G), IL-1α (H), and IL-1β (I). Results are from *n*=8-9 total mice per group per timepoint from *N*=2 independent experiments. (J) Representative images of unaltered whole lungs surgically collected from WT littermate and CCR2-DTR^+^ mice at Day 7 p.i. (K-N) Representative Masson’s trichrome stain of lung sections from WT littermate and CCR2-DTR^+^ mice comparing alveoli (I-J), scale bar = 100μM at 10X magnification, and bronchovascular bundles (K-L), scale bar = 25μM at 40X magnification. Airway (AW); Blood vessel (BV); Collagen fibers (black arrows); Fungal cells (yellow arrowheads). Data presented as mean ± SEM and analyzed using two-way ANOVA (A-I). *, *P* <0.05. **, *P* <0.01. ***, *P* <0.001. ****, *P* <0.0001.

**S3 Fig.** Dendritic cells do not have a role in the response to *C. gattii* infection. (A-B) Flow cytometry of dendritic cells from the lungs of MHCII^fl/fl^ controls (black line/white circles) and CCR2-Cre^+^ MHCII^fl/fl^ mice (orange filled line/orange circles) at Day 7 p.i., including a histogram of MHCII expression (A) and total numbers (B) of CD11b^+^ conventional DC (cDC), CD103^+^ cDC, and monocyte-derived DCs (moDCs). Data are *n*=6-10 total mice per group from *N=*2 independent experiments. (C) Kaplan-Meier survival curve of MHCII^fl/fl^ and CCR2-Cre^+^ MHCII^fl/fl^ mice, *n*=8-10 total mice per group from *N*=2 independent experiments. (D-F) Fungal burden was measured from lung (D), MLN (E), and brain (F) at Day 7 p.i. Data are *n*=7-9 total mice per group from *N*=2 independent experiments. (G-H) Flow cytometry of lung cells including CD11b^+^ cDC (G) and CD103^+^ cDC (H) from WT and *Csf2^-/-^* mice at Days 5 and 10 p.i. Data are *n*=5-8 total mice per group per time point from *N*=1-2 independent experiments. (I-J) Flow cytometry of lung cells from naive and infected WT littermates and infected CCR2-DTR^+^ mice at Days 3 and 7 p.i., including CD11b^+^ cDC (I) and CD103^+^ cDC (J). Results are from *n*=6 naive WT littermate mice from *N=*2 independent experiments and *n*=5 total infected mice per group per timepoint from *N*=1 experiment. Data presented as mean ± SEM and analyzed using two-way ANOVA (G-J), Mann-Whitney test (B, D-F), or Mantel-Cox test (C). ns, not significant. *, *P* <0.05. **, *P* <0.01. ***, *P* <0.001. ****, *P* <0.0001.

**S4 Fig.** Pulmonary Th1 and Th2 cytokines and chemokines in WT versus *Csf2*^-/-^ mice. (A-C) Levels of pulmonary cytokines in infected WT and *Csf2^-/-^* mice including Th1-associated and inflammatory cytokines (A), Th2-associated cytokines (B), and chemokines (C). Results are from *n*=5-10 total mice per group per timepoint from *N*=1-2 independent experiments. Data presented as mean ± SEM and analyzed using two-way ANOVA. *, *P* < 0.05. **, *P* < 0.01. ***, *P* < 0.001. ****, *P* < 0.0001.

**S5 Fig.** Pulmonary Th1 and Th2 cytokines and chemokines in WT littermate versus CCR2-DTR^+^ mice. (A-C) Levels of pulmonary cytokines in infected WT littermate and CCR2-DTR^+^ mice including Th1-associated and inflammatory cytokines (A), Th2-associated cytokines (B), and chemokines (C). Results are from *n*=5-10 total mice per group per timepoint from *N*=1-2 independent experiments. Data presented as mean ± SEM and analyzed using two-way ANOVA. *, *P* < 0.05. **, *P* < 0.01. ***, *P* < 0.001. ****, *P* < 0.0001.

**S6 Fig.** GM-CSF and monocyte-derived macrophages promote airway inflammation. (A-B) Representative Masson’s trichrome stained sections showing large airways, small airways, and terminal airways in the lungs of WT and *Csf2^-/-^* mice at Day 10 p.i. (A) and WT littermate and CCR2-DTR^+^ mice at Day 7 p.i. (B). Images in the left panels for each mouse strain are shown with scale bar = 100μM at 10X magnification. Black boxes represent magnified insets shown in the right panels for each mouse strain, with scale bar = 12μM at 60X magnification. Fungal cells (yellow arrowheads), goblet cells (green arrowheads); subepithelial layer (orange arrows).

**S7 Fig.** Gating strategy for macrophage, monocyte, and granulocyte cell populations in murine lungs by flow cytometry. Lung cells from a naive CCR2-DTR^+^ mouse were collected for flow cytometry analysis. Gating strategy for macrophages (CD45^+^CD64^+^MerTK^+^) including alveolar macrophages (CD11b^−^CD11c^hi^SiglecF^+^) and interstitial macrophages (CD11b^+^CD11c^+^SiglecF^−^); monocytes (CD45^+^CD64^+^MerTK^−^CD11c^−^SiglecF^-^CD11b^+^) including Ly6C^hi^ monocytes and Ly6C^lo^ monocytes; and granulocytes (CD45^+^CD64^−^MHCII^-−^CD11c^−^CD11b^+^SSC^hi^) including neutrophils (Ly6G^+^SiglecF^−^) and eosinophils (Ly6G^−^SiglecF^+^).

**S8 Fig.** Gating strategy for dendritic cell subsets in murine lungs by flow cytometry. Lung cells from a naïve CCR2-DTR^+^ mouse were collected for flow cytometry analysis. This strategy first excludes SiglecF^+^, MerTK^+^, CD3^+^, and CD19^+^ cells and then identifies DCs (CD45^+^CD3^−^CD19^−^MerTK^−^SiglecF^−^ MHCII^hi^ CD11c^+^) including CD11b^+^ cDCs (CD64^−^CD11b^+^CD103^−^), CD103^+^ cDCs (CD64^−^ CD11b^−^CD103^+^), and moDCs (CD64^+^CD11b^+^ CD103^−^).

**S9 Fig.** Gating strategy for MHCII expression by DCs in the lungs of CCR2-Cre^+^ MHCII^fl/fl^ mice. Lung cells from MHCII^fl/fl^ littermate controls were collected for flow cytometry analysis on Day 7 p.i. This strategy first excludes SiglecF^+^, MerTK^+^, CD3^+^, and CD19^+^ cells and then separates monocyte-derived and non-monocyte-derived cells by CD64 histogram. FSC-A^hi^ cells were selected for CD11b^+^ vs CD103^+^ gating. CD11b^+^ and CD103^+^ cDCs and moDCs were then determined by using a MHCII expression overlay to determine MHCII^+^ cell populations in the littermate control mice. These overlays were then applied to DC gates in CCR2-Cre^+^ MHCII^fl/fl^ samples.

**S1 Table.** Conjugated antibodies used for flow cytometry experiments.

## References

1. Heitman J, Kozel, T. R., Kwon-Chung, K. J., Perfect, J. R. and Casadevall A. Cryptococcus: from human pathogen to model yeast. 1 ed: ASM Press; 2010 November 12, 2010. 1–646 p.

2. Stephen C, Lester S, Black W, Fyfe M, Raverty S. Multispecies outbreak of cryptococcosis on southern Vancouver Island, British Columbia. Can Vet J. 2002;43(10):792–4. pmid:12395765.

3. Hoang LM, Maguire JA, Doyle P, Fyfe M, Roscoe DL. *Cryptococcus neoformans* infections at Vancouver Hospital and Health Sciences Centre (1997–2002): epidemiology, microbiology and histopathology. J Med Microbiol. 2004;53(9):935–40. pmid:15314203.

4. Casalini G, Giacomelli A, Antinori S. The WHO fungal priority pathogens list: a crucial reappraisal to review the prioritisation. Lancet Microbe. 2024;5(7):717–24.

5. Galanis E, Macdougall L, Kidd S, Morshed M. Epidemiology of *Cryptococcus gattii*, British Columbia, Canada, 1999-2007. Emerg Infect Dis. 2010;16(2):251–7. pmid:20113555.

6. Harris JR, Lockhart SR, Debess E, Marsden-Haug N, Goldoft M, Wohrle R, et al. *Cryptococcus gattii* in the United States: clinical aspects of infection with an emerging pathogen. Clin Infect Dis. 2011;53(12):1188–95. Epub 20111019. pmid:22016503.

7. Harris JR, Lockhart SR, Sondermeyer G, Vugia DJ, Crist MB, D’Angelo MT, et al. *Cryptococcus gattii* infections in multiple states outside the US Pacific Northwest. Emerg Infect Dis. 2013;19(10):1620–6. pmid:24050410.

8. WHO. WHO fungal priority pathogens list to guide research, development and public health action.: World Health Organization; 2022. Available from: https://www.who.int/publications/i/item/9789240060241

9. Rosen LB, Freeman AF, Yang LM, Jutivorakool K, Olivier KN, Angkasekwinai N, et al. Anti-GM-CSF autoantibodies in patients with cryptococcal meningitis. J Immunol. 2013;190(8):3959–66. pmid:23509356.

10. Jiang YK, Zhou LH, Cheng JH, Zhu JH, Luo Y, Li L, et al. Anti-GM-CSF autoantibodies predict outcome of cryptococcal meningitis in patients not infected with HIV: A cohort study. Clin Microbiol Infect. 2024;30(5):660–5. Epub 20240129. pmid:38295989.

11. Kuo PH, Wu UI, Pan YH, Wang JT, Wang YC, Sun HY, et al. Neutralizing anti-granulocyte-macrophage colony-Stimulating factor autoantibodies in patients with central nervous system and localized cryptococcosis: longitudinal follow-up and literature Review. Clin Infect Dis. 2022;75(2):278–87. pmid:34718451.

12. Wang S-Y, Lo Y-F, Shih H-P, Ho M-W, Yeh C-F, Peng J-J, et al. Cryptococcus gattii infection as the major clinical manifestation in patients with autoantibodies against granulocyte–macrophage colony-stimulating factor J Clin Immunol. 2022;42(8):1730–41. pmid:35947322.

13. Ataya A, Knight V, Carey BC, Lee E, Tarling EJ, Wang T. The role of GM-CSF autoantibodies in infection and autoimmune pulmonary alveolar proteinosis: a concise review. Front Immunol. 2021;12:752856. Epub 2021/12/10. pmid:34880857.

14. Punatar AD, Kusne S, Blair JE, Seville MT, Vikram HR. Opportunistic infections in patients with pulmonary alveolar proteinosis. J Infect. 2012;65(2):173–9. Epub 2012/04/10. pmid:22484272.

15. Rosen LB, Rocha Pereira N, Figueiredo C, Fiske LC, Ressner RA, Hong JC, et al. Nocardia-induced granulocyte macrophage colony-stimulating factor is neutralized by autoantibodies in disseminated/extrapulmonary nocardiosis. Clin Infect Dis. 2015;60(7):1017–25. Epub 20141203. pmid:25472947.

16. Becher B, Tugues S, Greter M. GM-CSF: From growth factor to central mediator of tissue inflammation Immunity. 2016;45(5):963–73. Epub 2016/11/17. pmid:27851925.

17. Chen Y, Li F, Hua M, Liang M, Song C. Role of GM-CSF in lung balance and disease. Front Immunol. 2023;14:1158859. Epub 2023/04/21. pmid:37081870.

18. Stanley E, Lieschke GJ, Grail D, Metcalf D, Hodgson G, Gall JA, et al. Granulocyte/macrophage colony-stimulating factor-deficient mice show no major perturbation of hematopoiesis but develop a characteristic pulmonary pathology. Proc Natl Acad Sci U S A. 1994;91(12):5592–6. Epub 1994/06/07. pmid:8202532.

19. Guilliams M, De Kleer I, Henri S, Post S, Vanhoutte L, De Prijck S, et al. Alveolar macrophages develop from fetal monocytes that differentiate into long-lived cells in the first week of life via GM-CSF. J Exp Med. 2013;210(10):1977–92. Epub 2013/09/18. pmid:24043763.

20. Misharin AV, Morales-Nebreda L, Reyfman PA, Cuda CM, Walter JM, McQuattie-Pimentel AC, et al. Monocyte-derived alveolar macrophages drive lung fibrosis and persist in the lung over the life span. J Exp Med. 2017;214(8):2387–404. Epub 2017/07/12. pmid:28694385.

21. Machiels B, Dourcy M, Xiao X, Javaux J, Mesnil C, Sabatel C, et al. A gammaherpesvirus provides protection against allergic asthma by inducing the replacement of resident alveolar macrophages with regulatory monocytes. Nat Immunol. 2017;18(12):1310–20. Epub 2017/10/17. pmid:29035391.

22. Aegerter H, Kulikauskaite J, Crotta S, Patel H, Kelly G, Hessel EM, et al. Influenza-induced monocyte-derived alveolar macrophages confer prolonged antibacterial protection. Nat Immunol. 2020;21(2):145–57. Epub 2020/01/15. pmid:31932810.

23. Evren E, Ringqvist E, Tripathi KP, Sleiers N, Rives IC, Alisjahbana A, et al. Distinct developmental pathways from blood monocytes generate human lung macrophage diversity. Immunity. 2021;54(2):259–75.e7.

24. Dal-Secco D, Wang J, Zeng Z, Kolaczkowska E, Wong CH, Petri B, et al. A dynamic spectrum of monocytes arising from the in situ reprogramming of CCR2+ monocytes at a site of sterile injury. J Exp Med. 2015;212(4):447–56. Epub 2015/03/25. pmid:25800956.

25. Mildner A, Schönheit J, Giladi A, David E, Lara-Astiaso D, Lorenzo-Vivas E, et al. Genomic Characterization of Murine Monocytes Reveals C/EBP&#x3b2; Transcription Factor Dependence of Ly6C^&#x2212;^ Cells. Immunity. 2017;46(5):849–62.e7.

26. Shibata Y, Berclaz PY, Chroneos ZC, Yoshida M, Whitsett JA, Trapnell BC. GM-CSF regulates alveolar macrophage differentiation and innate immunity in the lung through PU.1. Immunity. 2001;15(4):557–67. Epub 2001/10/24. pmid:11672538.

27. Chen G-H, Teitz-Tennenbaum S, Neal LM, Murdock BJ, Malachowski AN, Dils AJ, et al. Local GM-CSF–dependent differentiation and activation of pulmonary dendritic cells and macrophages protect against progressive cryptococcal lung infection in mice J Immunol. 2016;196(4):1810–21. pmid:26755822.

28. Osterholzer JJ, Chen GH, Olszewski MA, Curtis JL, Huffnagle GB, Toews GB. Accumulation of CD11b^+^ lung dendritic cells in response to fungal infection results from the CCR2-mediated recruitment and differentiation of Ly-6C^high^ monocytes. J Immunol. 2009;183(12):8044–53. pmid:19933856.

29. Gelman AE, Okazaki M, Sugimoto S, Li W, Kornfeld CG, Lai J, et al. CCR2 regulates monocyte recruitment as well as CD4 T1 allorecognition after lung transplantation. Am J Transplant. 2010;10(5):1189–99. pmid:20420631.

30. Liu X, Boyer MA, Holmgren AM, Shin S. Legionella-infected macrophages engage the alveolar epithelium to metabolically reprogram myeloid cells and promote antibacterial inflammation. Cell Host Microbe. 2020;28(5):683–98.e6.

31. Hohl TM, Rivera A, Lipuma L, Gallegos A, Shi C, Mack M, et al. Inflammatory monocytes facilitate adaptive CD4 T cell responses during respiratory fungal infection. Cell Host Microbe. 2009;6(5):470–81. pmid:19917501.

32. Bosteels C, Fierens K, De Prijck S, Van Moorleghem J, Vanheerswynghels M, De Wolf C, et al. CCR2- and Flt3-dependent inflammatory conventional type 2 dendritic cells are necessary for the induction of adaptive immunity by the human vaccine adjuvant system AS01. Front Immunol. 2020;11:606805. Epub 20210114. pmid:33519816.

33. Lewis Kanako L, Caton Michele L, Bogunovic M, Greter M, Grajkowska Lucja T, Ng D, et al. Notch2 receptor signaling controls functional differentiation of dendritic cells in the spleen and intestine. Immunity. 2011;35(5):780–91.

34. Greter M, Helft J, Chow A, Hashimoto D, Mortha A, Agudo-Cantero J, et al. GM-CSF controls nonlymphoid Tissue dendritic cell homeostasis but Is dispensable for the differentiation of inflammatory dendritic cells. Immunity. 2012;36(6):1031–46.

35. Voelz K, Johnston SA, Rutherford JC, May RC. Automated analysis of cryptococcal macrophage parasitism using GFP-tagged cryptococci. PLoS One. 2010;5(12):e15968. pmid:21209844.

36. Heung LJ, Hohl TM. Inflammatory monocytes are detrimental to the host immune response during acute infection with *Cryptococcus neoformans*. PLoS Patho. 2019;15(3):e1007627. Epub 2019/03/22. pmid:30897162.

37. Ballesteros I, Rubio-Ponce A, Genua M, Lusito E, Kwok I, Fernández-Calvo G, et al. Co-option of neutrophil fates by tissue environments. Cell. 2020;183(5):1282–97.e18.

38. MacDougall L, Kidd SE, Galanis E, Mak S, Leslie MJ, Cieslak PR, et al. Spread of *Cryptococcus gattii* in British Columbia, Canada, and detection in the Pacific Northwest, USA. Emerg Infect Dis. 2007;13(1):42–50. pmid:17370514.

39. MacDougall L, Fyfe M. Emergence of *Cryptococcus gattii* in a Novel Environment Provides Clues to Its Incubation Period. Journal of Clinical Microbiology. 2006;44(5):1851–2.

40. Kidd SE, Hagen F, Tscharke RL, Huynh M, Bartlett KH, Fyfe M, et al. A rare genotype of *Cryptococcus gattii* caused the cryptococcosis outbreak on Vancouver Island (British Columbia, Canada). Proceedings of the National Academy of Sciences. 2004;101(49):17258–63.

41. Hansakon A, Khampoongern R, Schiller L, Jeerawattanawart S, Angkasekwinai P. Effect of intranasal administration of granulocyte-macrophage colony-stimulating factor on pulmonary *Cryptococcus gattii* infection. Int Immunopharmacol. 2024;142:113259. pmid:39332096.

42. Oliveira-Brito PKM, de Campos GY, Guimarães JG, Serafim da Costa L, Silva de Moura E, Lazo-Chica JE, et al. Adjuvant curdlan contributes to immunization against *Cryptococcus gattii* infection in a mouse strain-specific manner. Vaccines. 2022;10(4):620. pmid:10.3390/vaccines10040620.

43. Chen GH, Curtis JL, Mody CH, Christensen PJ, Armstrong LR, Toews GB. Effect of granulocyte-macrophage colony-stimulating factor on rat alveolar macrophage anticryptococcal activity *in vitro*. J Immunol. 1994;152(2):724–34.

44. Chen G-H, Olszewski MA, McDonald RA, Wells JC, Paine R, Huffnagle GB, et al. Role of granulocyte macrophage colony-stimulating factor in host defense against pulmonary *Cryptococcus neoformans* infection during murine allergic bronchopulmonary mycosis Am J Pathol. 2007;170(3):1028–40. pmid:17322386.

45. Chiller T, Farrokhshad K, Brummer E, Stevens DA. Effect of granulocyte colony-stimulating factor and granulocyte-macrophage colony-stimulating factor on polymorphonuclear neutrophils, monocytes or monocyte-derived macrophages combined with voriconazole against *Cryptococcus neoformans*. Med Mycol J. 2002;40(1):21–6. pmid:11860010.

46. Strickland AB, Chen Y, Sun D, Shi M. Alternatively activated lung alveolar and interstitial macrophages promote fungal growth. iScience. 2023;26(5):106717.

47. Hansakon A, Mutthakalin P, Ngamskulrungroj P, Chayakulkeeree M, Angkasekwinai P. *Cryptococcus neoformans* and *Cryptococcus gattii* clinical isolates from Thailand display diverse phenotypic interactions with macrophages. Virulence. 2019;10(1):26–36. Epub 2018/12/07. pmid:30520685.

48. Piffer AC, Santos FMD, Thomé MP, Diehl C, Garcia AWA, Kinskovski UP, et al. Transcriptomic analysis reveals that mTOR pathway can be modulated in macrophage cells by the presence of cryptococcal cells. Genet Mol Biol. 2021;44(3):e20200390. Epub 2021/08/06. pmid:34352067.

49. Jackson DJ, Akuthota P, Roufosse F. Eosinophils and eosinophilic immune dysfunction in health and disease. Eur Respir Rev. 2022;31(163). Epub 2022/01/28. pmid:35082127.

50. Brightling CE, Monteiro W, Ward R, Parker D, Morgan MDL, Wardlaw AJ, et al. Sputum eosinophilia and short-term response to prednisolone in chronic obstructive pulmonary disease: a randomised controlled trial. Lancet. 2000;356(9240):1480-5.

51. Brightling CE, Symon FA, Birring SS, Bradding P, Wardlaw AJ, Pavord ID. Comparison of airway immunopathology of eosinophilic bronchitis and asthma. Thorax. 2003;58(6):528–32.

52. Varricchi G, Brightling CE, Grainge C, Lambrecht BN, Chanez P. Airway remodelling in asthma and the epithelium: on the edge of a new era. Eur Respir J. 2024;63(4). Epub 2024/04/13. pmid:38609094.

53. Zhou Y, Duan Q, Yang D. In vitro human cell-based models to study airway remodeling in asthma. Biomed Pharmacother. 2023;159:114218. Epub 2023/01/14. pmid:36638596.

54. Nobs SP, Kayhan M, Kopf M. GM-CSF intrinsically controls eosinophil accumulation in the setting of allergic airway inflammation. J Allergy Clin Immunol. 2019;143(4):1513–24.e2. pmid:30244025.

55. Voehringer D, van Rooijen N, Locksley RM. Eosinophils develop in distinct stages and are recruited to peripheral sites by alternatively activated macrophages. J Leukoc Biol. 2007;81(6):1434–44. Epub 2007/03/07. pmid:17339609.

56. Dougan M, Dranoff G, Dougan SK. GM-CSF, IL-3, and IL-5 Family of Cytokines: Regulators of Inflammation. Immunity. 2019;50(4):796–811. Epub 2019/04/18. pmid:30995500.

57. Oliveira MA, Lima GM, Shio MT, Leenen PJ, Abrahamsohn IA. Immature macrophages derived from mouse bone marrow produce large amounts of IL-12p40 after LPS stimulation. J Leukoc Biol. 2003;74(5):857–67. pmid:14595006.

58. Ling P, Gately MK, Gubler U, Stern AS, Lin P, Hollfelder K, et al. Human IL-12 p40 homodimer binds to the IL-12 receptor but does not mediate biologic activity. J Immunol. 1995;154(1):116–27. pmid:7527811.

59. Mattner F, Fischer S, Guckes S, Jin S, Kaulen H, Schmitt E, et al. The interleukin-12 subunit p40 specifically inhibits effects of the interleukin-12 heterodimer. Eur J Immunol. 1993;23(9):2202–8. Epub 1993/09/01. pmid:8103745.

60. Gillessen S, Carvajal D, Ling P, Podlaski FJ, Stremlo DL, Familletti PC, et al. Mouse interleukin-12 (IL-12) p40 homodimer: a potent IL-12 antagonist. Eur J Immunol. 1995;25(1):200–6. Epub 1995/01/01. pmid:7843232.

61. Heinzel FP, Hujer AM, Ahmed FN, Rerko RM. In vivo production and function of IL-12 p40 homodimers. J Immunol. 1997;158(9):4381–8. Epub 1997/05/01. pmid:9127002.

62. Nutku E, Zhuang Q, Soussi-Gounni A, Aris F, Mazer BD, Hamid Q. Functional expression of IL-12 receptor by human eosinophils: IL-12 promotes eosinophil apoptosis. J Immunol. 2001;167(2):1039–46. pmid:11441113.

63. Moreno SE, Alves-Filho JC, Alfaya TM, da Silva JS, Ferreira SH, Liew FY. IL-12, but not IL-18, is critical to neutrophil activation and resistance to polymicrobial sepsis induced by cecal ligation and puncture. J Immunol. 2006;177(5):3218–24. pmid:16920961.

64. McGeachy MJ, Cua DJ, Gaffen SL. The IL-17 Family of Cytokines in Health and Disease. Immunity. 2019;50(4):892–906. Epub 2019/04/18. pmid:30995505.

65. Murdock BJ, Falkowski NR, Shreiner AB, Sadighi Akha AA, McDonald RA, White ES, et al. Interleukin-17 drives pulmonary eosinophilia following repeated exposure to Aspergillus fumigatus conidia. Infect Immun. 2012;80(4):1424–36. Epub 20120117. pmid:22252873.

66. Malacco NLSdO, Rachid MA, Gurgel ILdS, Moura TR, Sucupira PHF, Sousa LPd, et al. Eosinophil-associated innate IL-17 response promotes aspergillus fumigatus lung pathology. Front Cell Infect Microbiol. 2019;Volume 8 - 2018.

67. Martin KR, Wong HL, Witko-Sarsat V, Wicks IP. G-CSF - A double edge sword in neutrophil mediated immunity. Semin Immunol. 2021;54:101516. Epub 2021/11/04. pmid:34728120.

68. Ueno K, Yanagihara N, Otani Y, Shimizu K, Kinjo Y, Miyazaki Y. Neutrophil-mediated antifungal activity against highly virulent *Cryptococcus gattii* strain R265. Med Mycol. 2019;57(8):1046–54. Epub 2019/01/23. pmid:30668754.

69. Mills KAM, Westermann F, Espinosa V, Rosiek E, Desai JV, Aufiero MA, et al. GM-CSF-mediated epithelial-immune cell cross-talk orchestrates pulmonary immunity to *Aspergillus fumigatus*. Sci Immunol. 2025;10(105):eadr0547.

70. Kasahara S, Jhingran A, Dhingra S, Salem A, Cramer RA, Hohl TM. Role of granulocyte-macrophage colony-stimulating factor signaling in regulating neutrophil antifungal activity and the oxidative burst during respiratory fungal challenge. J Infect Dis. 2016;213(8):1289–98.

71. Deepe GS, Jr., Gibbons R, Woodward E. Neutralization of endogenous granulocyte-macrophage colony-stimulating factor subverts the protective immune response to *Histoplasma capsulatum*. J Immunol. 1999;163(9):4985–93.

72. Hernández-Santos N, Wiesner DL, Fites JS, McDermott AJ, Warner T, Wüthrich M, et al. Lung epithelial cells coordinate innate lymphocytes and immunity against pulmonary fungal infection. Cell Host Microbe. 2018;23(4):511–22.e5.

73. Huang C, Wang Y, Li X, Ren L, Zhao J, Hu Y, et al. Clinical features of patients infected with 2019 novel coronavirus in Wuhan, China. Lancet. 2020;395(10223):497–506. pmid:31986264.

74. Thwaites RS, Sanchez Sevilla Uruchurtu A, Siggins MK, Liew F, Russell CD, Moore SC, et al. Inflammatory profiles across the spectrum of disease reveal a distinct role for GM-CSF in severe COVID-19. Sci Immunol. 2021;6(57):eabg9873.

75. De Luca G, Cavalli G, Campochiaro C, Della-Torre E, Angelillo P, Tomelleri A, et al. GM-CSF blockade with mavrilimumab in severe COVID-19 pneumonia and systemic hyperinflammation: a single-centre, prospective cohort study. Lancet Rheumatol. 2020;2(8):e465–e73.

76. Temesgen Z, Burger CD, Baker J, Polk C, Libertin CR, Kelley CF, et al. Lenzilumab in hospitalised patients with COVID-19 pneumonia (LIVE-AIR): a phase 3, randomised, placebo-controlled trial. Lancet Respir Med. 2022;10(3):237–46.

77. Arango-Franco CA, Rojas J, Firacative C, Migaud M, Agudelo CI, Franco JL, et al. Autoantibodies neutralizing GM-CSF in HIV-negative Colombian patients infected with *Cryptococcus gattii* and *C. neoformans*. J Clin Immunol. 2024;44(7):163.

78. Salvator H, Cheng A, Rosen LB, Williamson PR, Bennett JE, Kashyap A, et al. Neutralizing GM-CSF autoantibodies in pulmonary alveolar proteinosis, cryptococcal meningitis and severe nocardiosis. Respir Res. 2022;23(1):280. Epub 2022/10/12. pmid:36221098.

79. Vincent T, Plawecki M, Goulabchand R, Guilpain P, Eliaou JF. Emerging clinical phenotypes associated with anti-cytokine autoantibodies. Autoimmun Rev. 2015;14(6):528–35.

80. Cappellano G, Orilieri E, Woldetsadik AD, Boggio E, Soluri MF, Comi C, et al. Anti-cytokine autoantibodies in autoimmune diseases. Am J Clin Exp Immunol. 2012;1(2):136–46. Epub 20121115. pmid:23885320.

81. Jhingran A, Kasahara S, Hohl TM. Flow cytometry of lung and bronchoalveolar lavage fluid cells from mice challenged with Fluorescent Aspergillus Reporter (FLARE) Conidia. Bio Protoc. 2016;6(18). pmid:28691040.

82. Heung LJ, Hohl TM. DAP12 inhibits pulmonary immune responses to *Cryptococcus neoformans*. Infect Immun. 2016;84(6):1879–86. pmid:27068093.

